# Pharmacological inhibition of mTORC1 reduces neural death and damage volume after MCAO by modulating microglial reactivity

**DOI:** 10.1101/2023.12.31.571469

**Authors:** Mario Villa-González, Marina Rubio, Gerardo Martín-López, Paula R Mallavibarrena, Laura Vallés-Saiz, Denis Vivien, Francisco Wandosell, Maria José Pérez-Álvarez

## Abstract

Ischemic stroke is a sudden and acute disease characterized by neuronal death, glia activation, and a severe inflammatory process. Neuroinflammation is an early event after cerebral ischemia, with microglia playing a leading role. Microglial activation involves functional and morphological changes that drive a wide variety of phenotypes. In this context, deciphering the molecular mechanisms underlying such microglial activation is essential to devise strategies to protect neurons and maintain certain brain functions affected by early neuroinflammation after ischemia.

Here, we studied the role of mammalian target of rapamycin (mTOR) activity in the microglial response using a murine model of cerebral ischemia in the acute phase. We also determined the therapeutic relevance of the pharmacological administration of rapamycin, a mTOR inhibitor, before and after ischemic injury.

Our data show that rapamycin, administered before or after brain ischemia induction, reduced the volume of brain damage and neuronal loss by attenuating the microglial response. Therefore, our findings indicate that the pharmacological inhibition of mTORC1 in the acute phase of ischemia may provide an alternative strategy to reduce neuronal damage through attenuation of the associated neuroinflammation.

## Introduction

Neuroinflammation is a Central Nervous System (CNS) reactive response triggered by infection, trauma, autoimmune disorders or neurodegeneration (Glass et al., 2010). Recent evidence suggests that neuroinflammation aggravates damaged in neurological diseases, including ischemia (Jurcau & Simion, 2021).

Ischemic stroke is a sudden and acute disease characterized by a severe inflammatory process (Zietz et al., 2023). It represents the second cause of death and the first cause of long-term disability worldwide (J. Fan et al., 2023; Meschia & Brott, 2018). Brain ischemia is initiated by the total or partial blockage of a cerebral artery that generates a severe reduction of oxygen and glucose supply, promoting a cellular energy failure that trigger a sequence of pathological events that induces neuronal death (Dirnagl et al., 1999). The different degree of damage in the ischemic area allow to distinguish two different affected regions: the infarct zone or core, and the peri-infarct region. Core is the more affected region, characterized by a drastic reduction of blood flow, whereas peri-infarct region that surrounds the core, receives a sufficient supply of nutrients to maintain certain cellular functions (Ermine et al., 2021). It has been described that neuronal damage in infarct zone is almost irreversible given that is related with high rate of necrosis. By contrast, the less damaged peri-infarct zone is potentially recoverable with an early intervention (Mansfield et al., 2018). The only therapeutical approximation approved in acute phase is reperfusion, however, it does not reduce the consequent neurodegeneration (Villa-González et al., 2022). Therefore, it is necessary to find alternative treatments to reduce ischemic damage and improve the beneficial effects of reperfusion.

Neuroinflammation is an early event after cerebral ischemia, being microglia a leading player. It has been described in murine models that microglial response, that include their activation, begins 30 minutes after of the onset of ischemia (Rupalla et al., 1998). Microglial activation involves functional and morphological changes that drives a wide range of microglia phenotypes from pro-inflammatory to anti-inflammatory states (Paolicelli et al., 2022). During the acute phase of cerebral ischemia microglia release inflammatory factors that generate a proinflammatory environment that aggravates neuronal damage (Ge et al., 2023; Xia et al., 2022). Therefore, understanding the molecular mechanisms underlying this early microglial activation after ischemia could help to protect neurons and maintain brain function.

One of the main players controlling microglial functional/phenotypic changes is the PI3K/Akt/mTOR pathway (Chu et al., 2021). mTOR is a ubiquitous serine-threonine protein kinase, highly conserved, formed by two multiprotein complexes: mTORC1 and mTORC2. Both mTOR complexes (mTORCs) coordinate a wide range of anabolic and catabolic responses, so it is considered a pivotal player in the cellular homeostasis (Liu & Sabatini, 2020; Villa-González et al., 2022). Consequently, mTOR signalling dysregulation or dysfunction significantly affect CNS integrity (Karalis & Bateup, 2021). Several *in vivo* studies of brain ischemia have reported a decrease in mTORCs activity after ischemic damage, accompanied by neuronal death and neurological deficits (Liu et al., 2019; Mateos et al., 2016; Pérez-Álvarez et al., 2012). The increase in mTORCs activity in ischemia promotes neuronal survival and diminishes ischemic damage (Pan et al., 2022; Perez-Alvarez et al., 2015).

However, inhibition of mTORCs activity after or before ischemia using *in vivo* models has shown controversial results. Some reports have demonstrated detrimental effects after mTORCs inhibition while others not (Chauhan et al., 2015; Chi et al., 2021; Li et al., 2016; Liu et al., 2019; Yang et al., 2015). The involvement of mTOR along different factors such as the timing of the study, the degree of damage or the response of each cell type to ischemic injury, could explain these results. Thus, it is necessary to better understand the role of mTORCs in all backgrounds to understand these controversial reports.

In this study, we have evaluated the role of mTOR activity after cerebral ischemia focusing on microglial response during the acute phase, by using the mTOR inhibitor rapamycin. Our data demonstrate that rapamycin administration both before and after MCAO, reduces the ischemic volume and neuronal loss and attenuates microglial response. Specifically, rapamycin reduces the levels of the pro-inflammatory microglial phenotype and promotes neuroprotection. These results suggest that the pharmacological inhibition of mTORC1, during the acute phase of ischemic stroke could be an alternative protective strategy to reduce neuronal damage through the attenuation

## Material and methods

### Animals and treatments

Swiss mice (12 weeks old, weight 35-45 g) were housed in the Animal Facility of U1237 (GIP Cyceron) with access to food and water *ad libitum* under a 12-hour light/dark cycle in a temperature-controlled environment. All care and experiments were performed following the ARRIVE guidelines (www.nc3rs.org.uk), including blind analyses of the samples. A stock solution of rapamycin at a concentration of 20 mg/mL in ethanol was prepared. Rapamycin was diluted in 5% polyethylene glycol-400 (PEG400, Fluka) and 5% Tween®80 (Sigma-Aldrich) in phosphate-buffered saline (PBS) and administered intraperitoneally at 20 mg/kg 48 h before the induction of MCAO (MCAO+R_pre_) or 20 min after (MCAO+R_post_).

We used a total of 57 male mice, which were randomly distributed into the following experimental groups (Figure Supplementary 1A): Sham+Vehicle (V), n=10; Sham+Rapamycin (R), n=10; MCAO+V, n=14; MCAO+R_pre,_ n=12; and MCAO+R_post_, n=11. We used 28 and 29 mice for western blot (WB) analysis and immunohistochemical analysis, respectively.

### Thromboembolic stroke model

Middle Cerebral Artery Occlusion (MCAO) by intravascular thrombin injection (Orset et al., 2007) was used. Animals were anesthetized with a mixture of 5% isoflurane, 1.5% O_2_, and 1.5% NO_2_ and were maintained with 2–3% isoflurane during the surgical procedure according to the needs of each animal. Once the animal had been fixed in the stereotaxic frame, a craniotomy was performed on the temporal bone. Dura mater and meninges were removed until the middle cerebral artery (MCA) was isolated. A glass micropipette was then introduced into the lumen of the base of the branching MCA and 1U of purified murine ɑ-thrombin (Enzyme Research Labs) was pneumatically introduced to induce clot formation. After 10 min, the micropipette was removed. Cerebral blood flow was monitored using a laser Doppler equipped with a fibre optic probe (Oxford Optronic) both before and up to 20 min after MCAO to verify and discard early spontaneous recanalization.

### Magnetic Resonance Imaging (MRI)

MRI using a Pharma Scan 7T (Bruker). was performed on all the animals 24 h after surgery. Sequence acquisition was carried out using a Pharma Scan 7T (Bruker). Images were acquired using a TE/TR 33 ms/2500 ms multi-slice sequence. In addition, a series of T2-weighted sequences and the angiogram were obtained to monitor MCA recanalization. Lesion size was determined using Image J software. Animals that showed a lesion < 5 mm^3^ 24 h post-MCAO were discarded.

### Western Blot (WB)

Soluble protein extracts from the parietal cortex of sham and operated mice were used for WB analysis. Mice under anaesthesia were subjected to intracardiac perfusion with PBS 7.4 pH. The damaged cortex was removed and maintained at −80°C until the subsequent analysis. Ice-cold lysis buffer [50 mM Tris 8.0 pH; 100 mM NaCl, 10 mM NaF, 5 mM EDTA, 1% Triton X-100, 1 μM okadaic acid, 2 mM sodium orthovanadate and protease inhibitors (Roche, #1697498)] was used to homogenize the brain samples. Homogenates were kept in ice for 15 min and centrifuged at 12,000 *g* for 10 min at 4°C. Protein concentration was measured using the DC Protein Assay (Bio-Rad), following the manufacturer’s protocol. Proteins were resolved on SDS-polyacrylamide gel, transferred to a 0.2 µm nitrocellulose blotting membrane (Amersham Protan), and blocked with 5% milk powder+0.1 % Tween in PBS. Membranes were incubated overnight at 4°C with primary antibodies diluted in 5% BSA+0.1% Tween in PBS (see Table 1). They were then washed with PBS and incubated for 1 h at room temperature with the appropriate secondary antibody: goat anti-Rabbit IgG-HRP (Southern Biotech, #4030-05) or goat anti-Mouse IgG-HRP (Southern Biotech, #1030-05) 1:5000. Immunoreactivity was detected using Clarity Western ECL substrate (Bio-Rad, #170-5061). β-Actin was used as an internal control. The relative expression levels of proteins were measured using ImageJ software (ImageJ, Fiji).

**Table 1.**
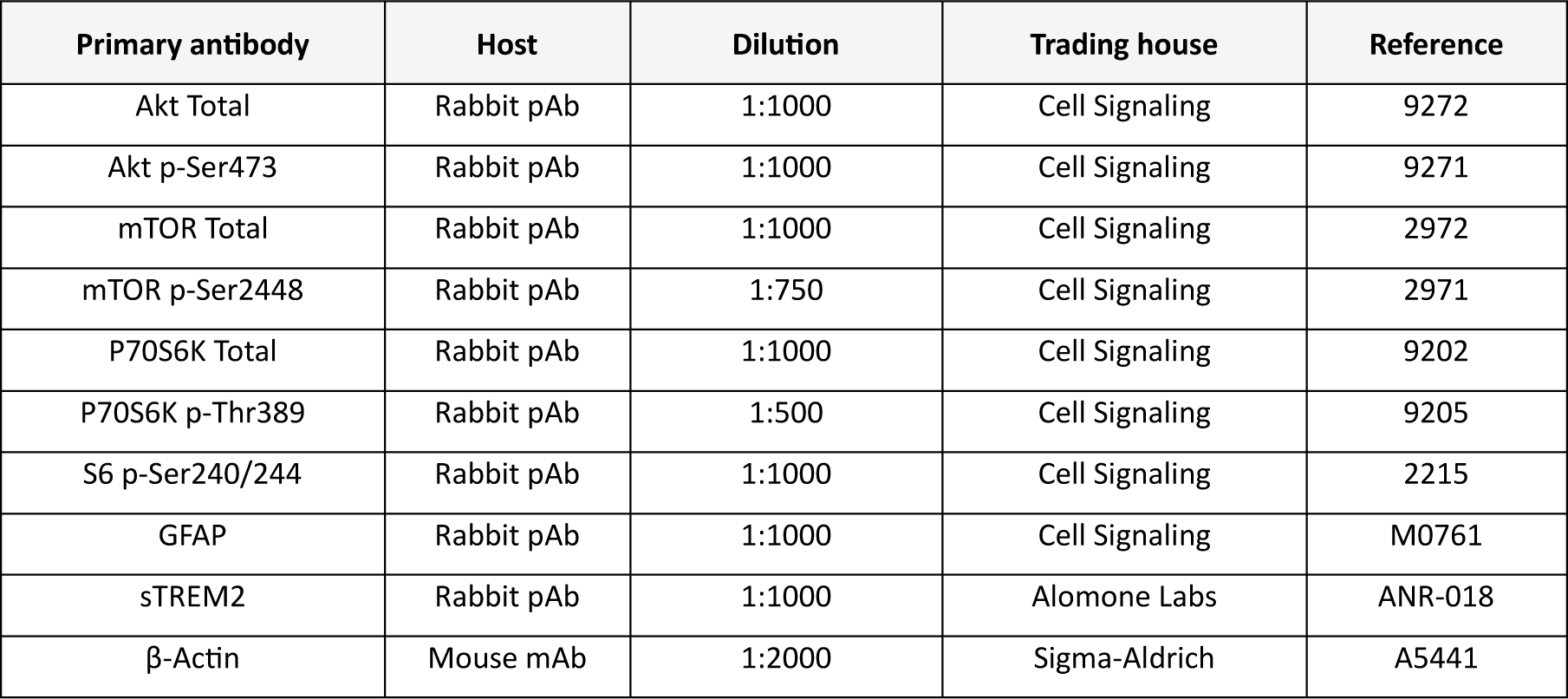
Primary antibodies used by Western Blot analysis.

| Primary antibody | Host | Dilution | Trading house | Reference |
| --- | --- | --- | --- | --- |
| Akt Total | Rabbit pAb | 1:1000 | Cell Signaling | 9272 |
| Akt p-Ser473 | Rabbit pAb | 1:1000 | Cell Signaling | 9271 |
| mTOR Total | Rabbit pAb | 1:1000 | Cell Signaling | 2972 |
| mTOR p-Ser2448 | Rabbit pAb | 1:750 | Cell Signaling | 2971 |
| P70S6K Total | Rabbit pAb | 1:1000 | Cell Signaling | 9202 |
| P70S6K p-Thr389 | Rabbit pAb | 1:500 | Cell Signaling | 9205 |
| S6 p-Ser240/244 | Rabbit pAb | 1:1000 | Cell Signaling | 2215 |
| GFAP | Rabbit pAb | 1:1000 | Cell Signaling | M0761 |
| sTREM2 | Rabbit pAb | 1:1000 | Alomone Labs | ANR-018 |
| β-Actin | Mouse mAb | 1:2000 | Sigma-Aldrich | A5441 |

### Tissue preparation

Mice were sacrificed by intracardiac perfusion with 4% paraformaldehyde in PBS (4% PFA) under 5% isoflurane 24 h after MCAO induction. Brains were immediately removed and fixed by immersion in 4% PFA for 24 h at 4°C. They were then cryoprotected using sucrose solution and finally embedded in Tissue-Tek medium (Sakura Finetek). Brain sections were cut into 10 µm thick coronal slices from Bregma +1 to −2 mm (Franklin, 2012).

### Immunohistochemistry and immunofluorescence

For immunohistochemical staining, brain sections were incubated in blocking solution (0.1% Triton X-100, 1% BSA, 1% Horse Serum in PBS) for 1 h and incubated overnight at 4°C with the specific primary antibody: mouse anti-NeuN 1:200 (Millipore, MAB377), rabbit anti-GFAP 1:1000 (Dako, M0761) or rabbit anti-Iba-1 1:500 (Wako, 019-19741) diluted in blocking solution. Endogenous peroxidase activity was inhibited with 0.1% H_2_O_2_ in PBS. After washing, samples were incubated with the appropriate secondary antibody using the ABC kit (Vectastain® ABC, Vector Laboratories, USA), following the manufacturer’s instructions. Secondary antibodies were visualized using diaminobenzidine (DAB). Coverslips were mounted with Depex (Panreac) and images were captured using an Olympus BX51 microscope with an Olympus camera DP-70 (Olympus).

For immunofluorescence staining, samples were pretreated for 30 min with 0.1 M Glycine 8.5 pH. They were then blocked for 1 h and incubated overnight at 4°C with guinea pig anti-Iba-1 1:750 (LabClinics, #HS-234004), rabbit anti-S6 p-Ser240/244 1:200 (Cell Signaling, #2215) or rabbit anti-sTREM2 1:300 (Alomone Labs, #ANR018) diluted in blocking solution. The following day, samples were washed in PBS and incubated with the appropriate secondary antibody: anti-rabbit Alexa 488 and anti-guinea pig Alexa 555, both 1:1000 (AlexaFluor, Thermo-Fisher). After washing, nuclei were stained with DAPI 1:5000 (Calbiochem) for 5 min. Sections were mounted with Fluoromount-G (Southern Biotechnology Associates).

Images were captured using a confocal laser scanning microscope LSM900 coupled to a vertical Axio Imager 2 microscope (Zeiss). Sequential optical sections (1 μm) were acquired in Z-stacks. We used a minimum of four brain slices from each mouse. Images from damaged regions (infarct and peri-infarct) were captured at 20X magnification. The number of positive cells was expressed as the mean of the total cell number counted *per* mm^2^ and it was calculated using the cell counter plugin with Fiji software. In brief, the image contrast was enhanced and then a threshold was established to eliminate the background.

### Analysis of microglial morphology

Microglial morphology was studied on Iba-1-stained immunofluorescence images from different non-consecutive sections from Bregma +1.2 to 0.5 mm taken at 20X magnification (Franklin, 2012). We used cell area and number of branch points as parameters for this analysis (Leyh et al., 2021). All cells present in the study field were analysed, using a minimum of 4 non-consecutive slices per mouse. In brief, the image contrast was enhanced and then a threshold was established to eliminate the background.

### Statistical analysis

All statistical analyses were performed with GraphPad Prism 8.0 (GraphPad Prism Software®). Data are represented as mean ± SEM (standard error of the mean). First, normal distribution of data was checked using the Shapiro-Wilk test. Two-way analysis of variance (ANOVA), followed by post hoc multiple comparisons using Bonferroni correction, was performed to compare more than two experimental groups. The unpaired t-test was used to compare two groups. All statistical tests were performed in a two-tailed manner. Data were considering statistically significant at p≤0.05 = *, p≤0.01 = **, p≤0.001 = ***.

## Results

### Rapamycin reduces the volume of ischemic lesions

The MRI analysis after MCAO induction showed that rapamycin administration significantly reduced lesion volume. MCAO induced a lesion of 17.4±1.7 mm^3^ and rapamycin-treated ischemic mice showed a decrease in lesion volume compared to untreated counterparts (MCAO+R_pre_ 11.4±1.6 mm^3^, p≤0.05; MCAO+R_post_ 9.6±1.5 mm^3^, p≤0.01 vs. MCAO). No differences were observed regarding the timing of rapamycin administration (Figure 1A-B).

**Figure 1.**
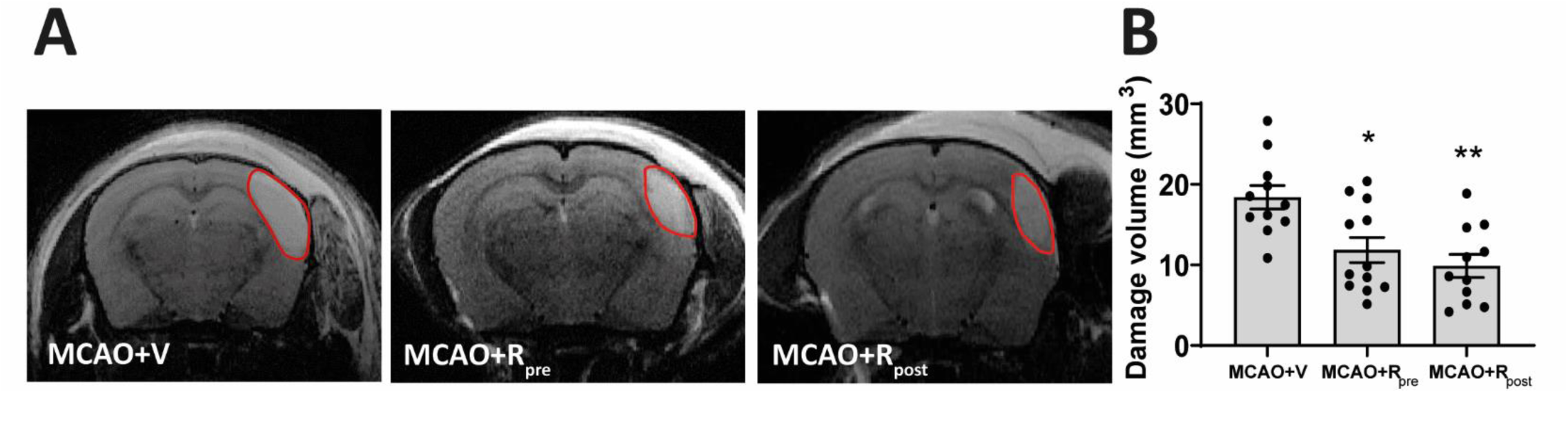
Rapamycin reduces volume of ischemic lesion. A) Representative MRI images showing the lesion volume after 24 hours of MCAO induction. The red line delineates the lesioned area. B) Graphical quantification of lesion volumes. Data represent means ± SEM (Ordinary one-way ANOVA *p ≤ 0.05, ** p≤0.01). MCAO+V n=11; MCAO+R_pre_ n=12; MCAO+R_post_ n=11 (V=vehicle; R=rapamycin).

### Rapamycin decreases mTORC1 activity in microglia after MCAO

Next, we analysed mTORC activity in brain tissue. We quantified the total amount and the phosphorylated levels of the main players of the mTORC1 pathway (see Figure 2A) to infer the activity levels of mTORC1 and mTORC2. To this end, we assessed the levels of Akt p-Ser473, which indicates mTORC2 activity, the levels of mTOR p-Ser2448 itself and P70S6K p-Thr389, the principal target of mTORC1 (Perez-Alvarez et al., 2018).

**Figure 2.**
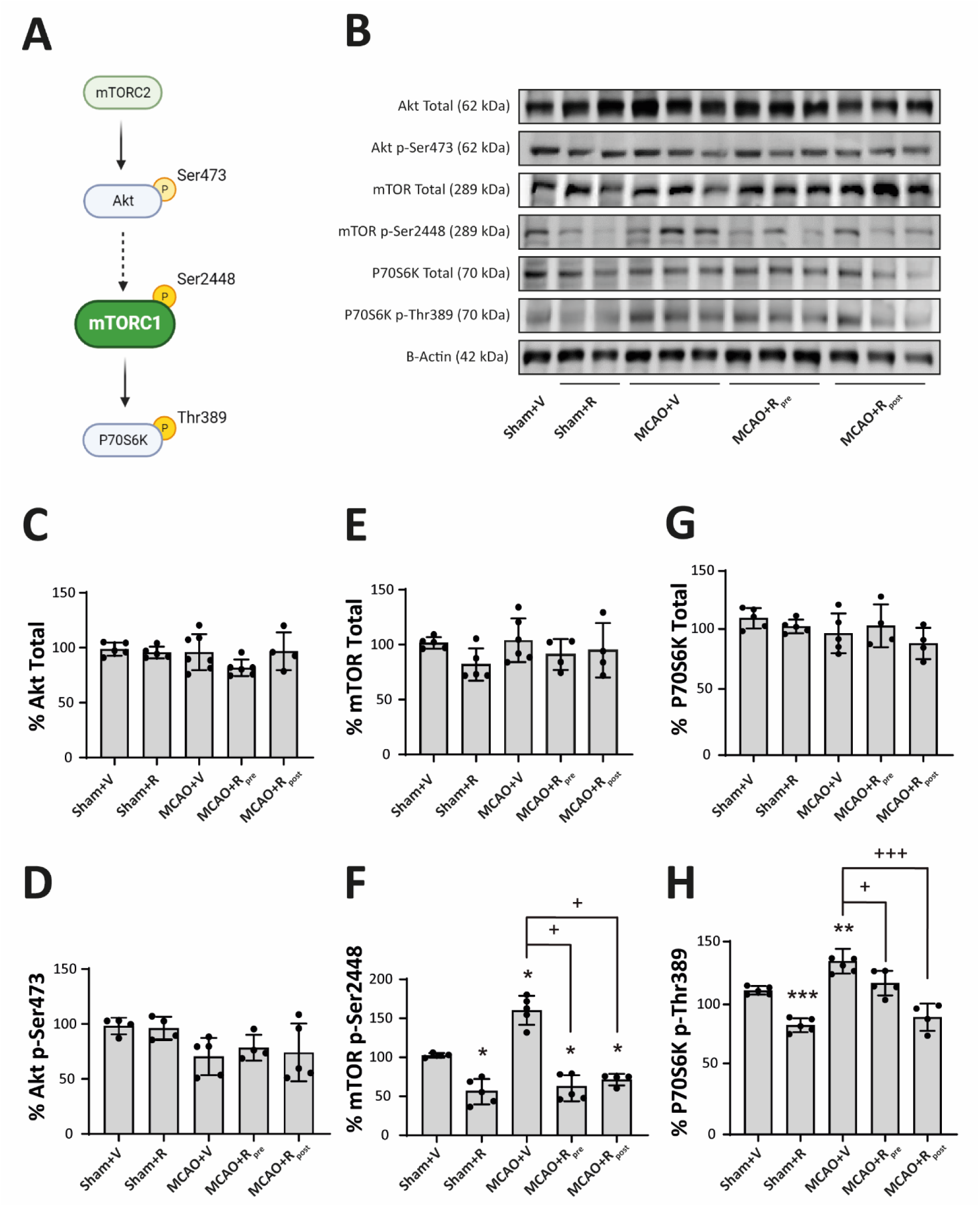
Rapamycin decreases mTORC1 activity in tissue after MCAO. A) Schematic representation of the mTORC1 pathway. B) Representative Western Blot images of the analysed extracts proteins from mice cerebral cortex. Immunodetections were performed using antibodies against total Akt, Akt p-Ser473, total mTOR, mTOR p-Ser2448, total P70S6K, P70S6K p-Thr389 and B-Actin, as a loading control. C-H) Graphical quantification of immunodetection by Western Blot. Data are normalized against B-Actin and expressed as the percentage of variation versus sham. Graph values represent means ± SEM (Ordinary one-way ANOVA *p ≤ 0.05; **p ≤ 0.01; ***p ≤ 0.001. Unparied t-test ^+^p ≤ 0.05; ^++^p ≤ 0.01; ^+++^p ≤ 0.001). Sham+V n=4-5, Sham+R n=4-5, MCAO+V n=5; MCAO+Rpre n=4-5 and MCAO+R_post_ n=4-5 (V=vehicle; R=rapamycin). * vs Sham and ^+^ vs MCAO+V group.

No variations in the levels of total Akt (Figure 2B-C), Akt p-Ser473 (Figure 2B-D), total mTOR (Figure 2B-E), or total P70S6K p-Thr389 (Figure 2B-G) were observed in any of the experimental groups. Rapamycin-treated sham mice showed a significant reduction in phosphorylation levels of mTOR p-Ser2448 (55.9±7.23%, p≤0.001) (Figure 2B-F) and P70S6K p-Thr389 (68.2±3.0%, p≤0.05) (Figure 2B-H) compared to untreated counterparts. These data confirm that rapamycin administration downregulates mTORC1 but not mTORC2 activity in the brain.

The MCAO group showed an increase in the levels of both phospho-epitopes compared to sham mice (p-mTOR: 55.9±7.3%, p≤0.05; p-P70S6K: 68.2±3%, p≤0.01) (Figure 2B-F and 2B-H). Rapamycin administration pre- and post-MCAO triggered a significant reduction in p-mTOR (MCAO+R_pre_: 103.3±7.2%, p≤0.05; MCAO+R_post_: 71.4±3.7%, p≤0.01) and p-P70S6K (MCAO+R_pre_: 109.4±5.4%, p≤0.05; MCAO+R_post_: 76±6.7%, p≤0.01) levels compared to the untreated ischemic group (Figure 2B-F and 2B-H). No differences were observed in mTOR phosphorylation levels regarding the timing of drug administration (Figure 2B-F). However, P70S6K p-Thr389 levels showed a large decrease when rapamycin was administered post-surgery compared to pre-surgery (76±6.7%, p≤0.01) (Figure 2B-H).

Next, we sought to elucidate the microglial contribution to increased mTORC1-activity after stroke in brain lysates. To this end, we examined the phosphorylation levels of S6 ribosomal protein, a direct target of mTORC1/P70S6K (Meyuhas, 2015), to infer mTORC1 activity in these cells by double immunofluorescence using p-S6 and Iba-1 antibodies (Figure 3A). p-S6^+^/Iba-1^+^ cells were absent in sham mice but present in the MCAO groups. The mice treated with rapamycin, pre- and post-MCAO, showed a decrease in the number of p-S6^+^/Iba-1^+^ cells compared to untreated ischemic mice. No differences were observed between pre- and post-rapamycin administration (Figure 3A). WB analyses indicated an increment in p-S6 levels after MCAO compared to sham mice (182.7±13.9%, p≤0.001) (Figure 3B-C). Rapamycin reduced p-S6 levels in treated sham mice compared to untreated ones (39.9±2.9%, p≤0.001) (Figure 3B-C). Similarly, both rapamycin-treated ischemic mouse groups showed a significant reduction in p-S6 levels compared to untreated ischemic (MCAO+R_pre_: 62.2%±2.8, p≤0.001; MCAO+R_post_: 43.1±4.4%, p≤0.001), without no differences between the drug administration times (Figure 3B-C). These data suggest that the mTORC1 activation observed by WB could be related to an increase in the activity of the PI3K/Akt/mTORC1 pathway in microglia.

**Figure 3.**
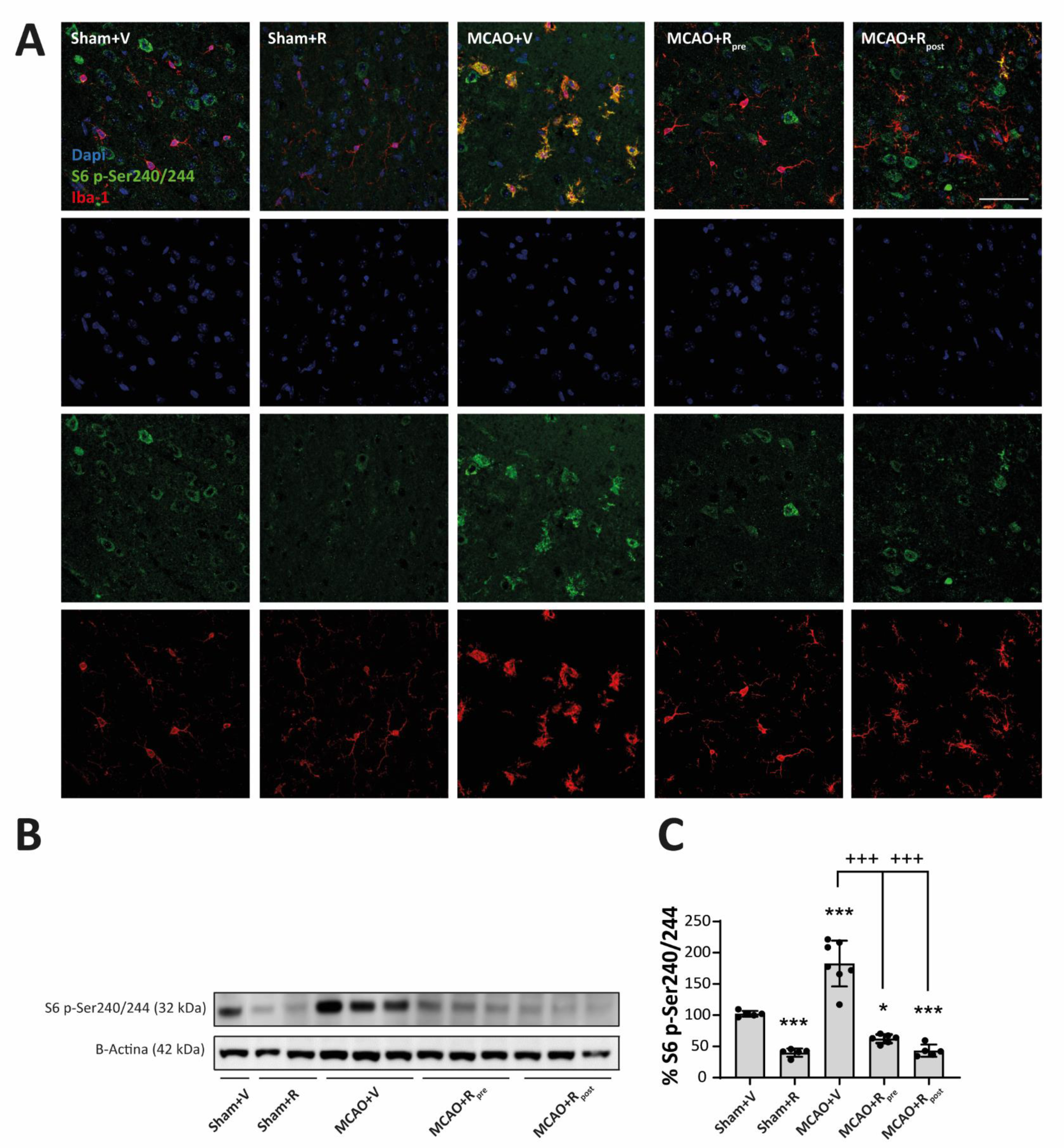
Rapamycin reduce mTORC1 activity in microglia after MCAO. A) Representative confocal immunofluorescence images. Staining was performed using Dapi (blue), S6 p-Ser240/244 (green) and Iba-1 (red) antibodies. Scale bar: 50 µm. B) Representative Western Blot images of the analysed extracts proteins from mice cerebral cortex. Immunodetections were performed using antibodies against S6 p-Ser240/244 and B-Actin, as a loading control. C) Graphical quantification of immunodetection by Western Blot. Data are normalized against B-Actin and expressed as the percentage of variation versus sham. Graph values represent means ± SEM (Ordinary one-way ANOVA *p ≤ 0.05; **p ≤ 0.01; ***p ≤ 0.001. Unparied t-test ^+^p ≤ 0.05; ^++^p ≤ 0.01; ^+++^p ≤ 0.001). Sham+V n=5, Sham+R n=5, MCAO+V n=7; MCAO+Rpre n=6 and MCAO+R_post_ n=5 (V=vehicle; R=rapamycin). * vs Sham+V and ^+^ vs MCAO+V group.

### Rapamycin reduces neuronal loss after MCAO

We studied the impact of mTORC1 inhibition pre- and post-MCAO on neuronal survival. To this end, we performed an immunohistochemistry assay using the NeuN antibody to quantify the number of NeuN^+^ cells in the damaged area (see Supplementary Figure 1B). Rapamycin treatment did not alter the number of NeuN^+^ cells in sham mice. MCAO reduced the total number of NeuN^+^ cells in the whole damaged area compared to sham mice (490±26 cells/mm^2^, p≤0.001) (Figure 4A-B). Regional analysis showed a greater decrease in the number of NeuN^+^ cells in the infarct zone (193±19 cells/mm^2^, p≤0.001) than in the peri-infarct region (297±19 cells/mm^2^, p≤0.001) (Figure 4A-C). MCAO mice treated with rapamycin showed a significant reduction in the loss of neurons throughout the damaged area compared to untreated ischemic, regardless of the time of drug administration (MCAO+R_pre_: 672±41 cells/mm^2^, p≤0.05; MCAO+R_post_: 639±34 cells/mm^2^, p≤0.01) (Figure 4A-B). In both rapamycin-treated ischemic groups, a further reduction in the number of NeuN^+^ cells in the infarct zone were observed (Figure 4A-C). No differences were found between the times of drug administration (Figure 4B-C). The data obtained from the WB analysis confirmed the histological results. MCAO significantly reduced NeuN levels compared to untreated sham mice (39.6±3.7%, p≤0.001), and rapamycin-treated ischemic mice showed an increase in total NeuN levels compared to untreated ischemic counterparts, both pre-(67.8±3.6%, p≤0.001) and post (60.5±1.9%, p≤0.001) ischemic injury. No differences were observed between the times of rapamycin administration (Figure 4D-E). These results suggest that mTORC1 inhibition prevents neuronal loss after ischemia.

**Figure 4.**
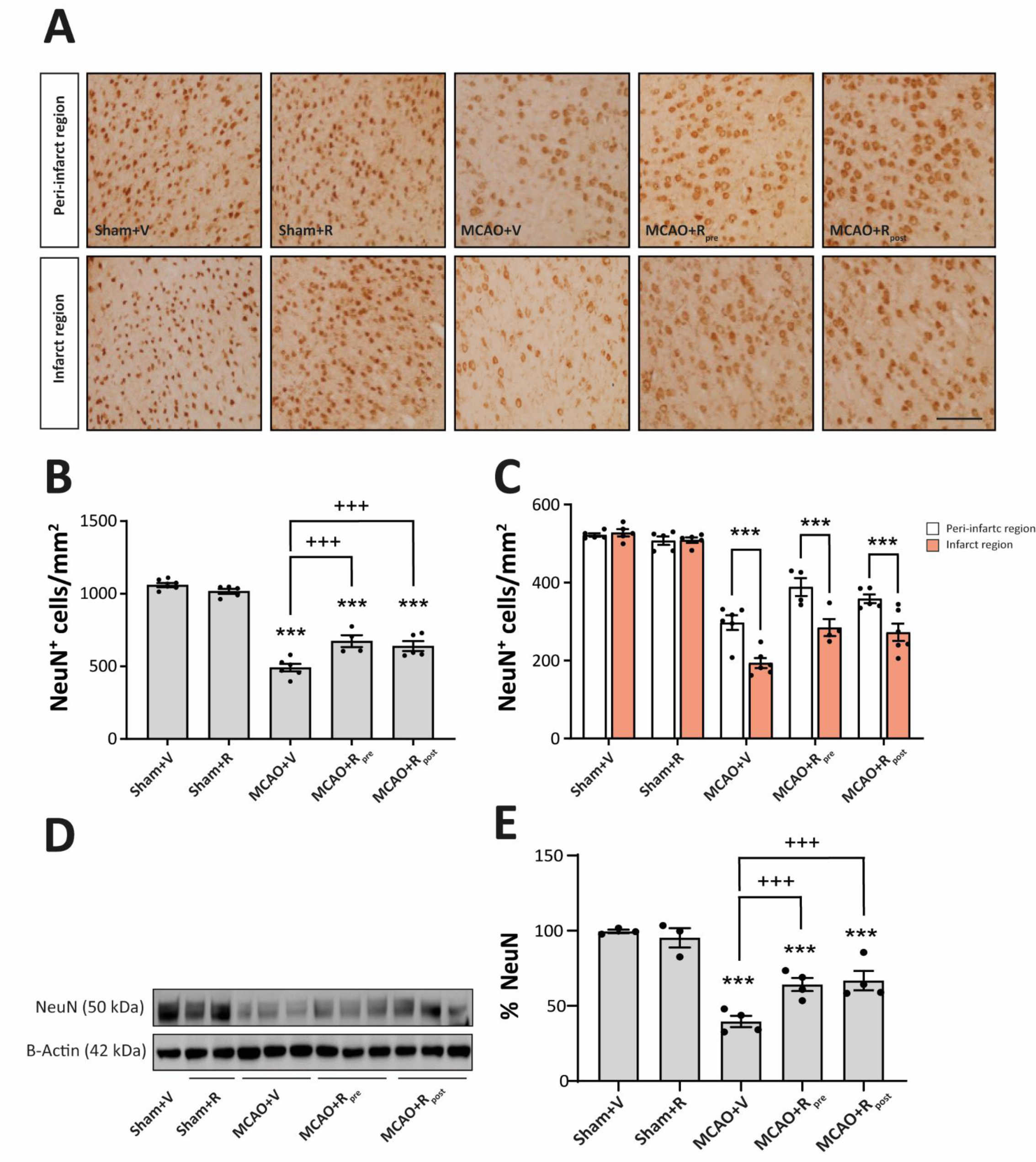
Rapamycin reduces neuronal loss after MCAO. A) Representative immunohistochemical images from cortex infarcted area and cortex peri-infarct region (coronal section, 10 µm). Staining was performed used NeuN antibody. Scale bar: 25 µm. B) Graphical quantification of NeuN^+^ cell counting across the entire damaged cortical region. C) Graphical quantification of NeuN^+^ cell counting, distinguishing between the two damaged regions (white bars represent peri-infarct region whereas red bars represent infarcted area). D) Representative Western Blot images of the analysed extracts proteins from mice cerebral cortex. Immunodetections were performed using antibodies against NeuN and B-Actin, as a loading control. E) Graphical quantification of immunodetection by Western Blot. Data are normalized against B-Actin and expressed as the percentage of variation versus sham. Graph values represent means ± SEM (Ordinary one/two-way ANOVA *p ≤ 0.05; **p ≤ 0.01; ***p ≤ 0.001. Unparied t-test ^+^p ≤ 0.05; ^++^p ≤ 0.01; ^+++^p ≤ 0.001). Sham+V n=5, Sham+R n=5, MCAO+V n=7; MCAO+Rpre n=6 and MCAO+R_post_ n=5 (V=vehicle; R=rapamycin). * vs Sham+V and ^+^ vs MCAO+V group.

### mTORC1 inhibition drives changes in microglial density and morphology after MCAO

Next, we performed immunohistochemistry on brain slices using the Iba-1 antibody to address the effect of mTORC1 inhibition specifically in microglia after MCAO. We used two distinct parameters to analyse the microglial response, namely the total number of Iba-1^+^ cells and the total stained area. In sham groups, rapamycin treatment did not affect either parameter (Figure 5B-C). In contrast, MCAO significantly increased both (762±42 cells/mm^2^, p≤0.01) (400.3±9.8%, p≤0.001) compared to untreated sham mice (Figure 5B-C). Rapamycin-treated ischemic mice showed a reduction in the number of Iba-1^+^ cells (MCAO+R_pre_: 560±30 cells/mm^2^, p≤0.001; MCAO+R_post_: 540±27 cells/mm^2^, p≤0.001) and the stained area (MCAO+R_pre_: 256.3±9.3%, p≤0.01; MCAO+R_post_: 277.3±8.4%, p≤0.001) compared to untreated ischemic counterparts and no differences were observed between the times of rapamycin administration (Figure 5B-C). No significant differences were observed between the damaged area of the infarct zone and peri-infarct region in any of the parameters analysed (data not shown).

**Figure 5.**
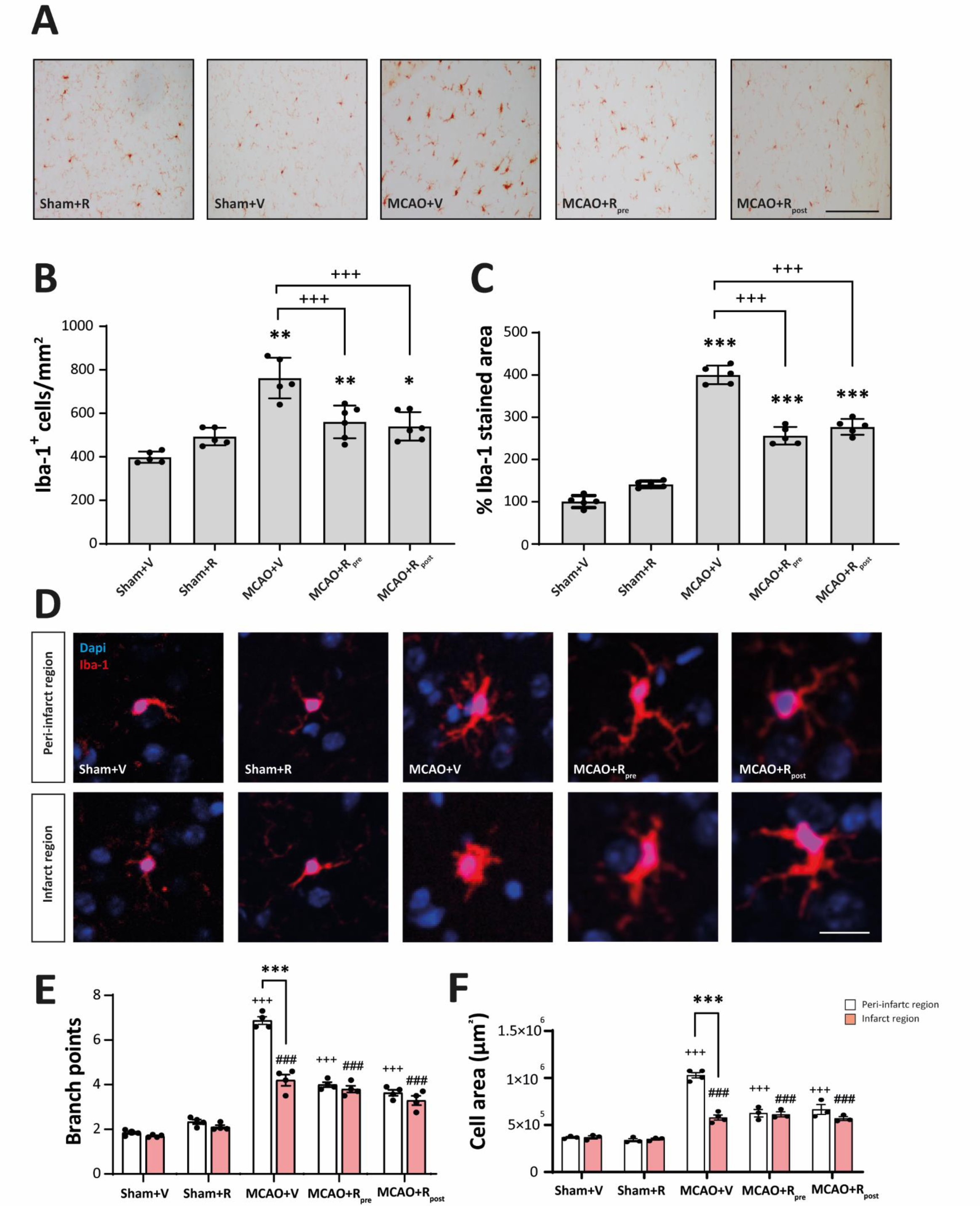
mTORC1 inhibition in microglia drives changes in their density and morphology after MCAO. A) Representative immunohistochemical images from the entire dagame area (coronal section, 10 µm). Staining was performed used Iba-1 antibody. Scale bar: 25 µm. B) Graphical representation of Iba-1^+^ cell counting and C) graphical representation of Iba-1^+^ stained damage area. D) Representative confocal immunofluorescence images from peri-infarct and infarct area. Staining was performed using Dapi (blue) and Iba-1 (red) antibody. Scale bar: 5 µm. E) Graphical representation of the quantification of branching points and the F) area occupied by each Iba-1^+^ cell in both damaged regions (white bars represent peri-infarct region whereas red bars represent infarcted area). Ordinary one/two-way ANOVA *p ≤ 0.05; **p ≤ 0.01; ***p ≤ 0.001. Unparied t-test ^+^p ≤ 0.05; ^++^p ≤ 0.01; ^+++^p ≤ 0.001). Sham+V n=5, Sham+R n=5, MCAO+V n=7; MCAO+Rpre n=6 and MCAO+R_post_ n=5 (V=vehicle; R=rapamycin). ^+^ vs Sham+V (peri-infarct region), ^#^ vs Sham+V (infarct region) and * vs same experimental group.

Microglial activation is a highly dynamic process that involves changes in cellular morphology. These variations have been associated with the extent and duration of ischemic damage (Hernández et al., 2021). Thus, we examined the effect of mTORC1 inhibition on the number of branch points and cell area of microglia (Leyh et al., 2021), addressing the infarct zone and peri-infarct region separately. Rapamycin administration in sham mice did not cause any variations in the number of branch points or cell area in either of the damaged regions (Figure 5E-F). In contrast, mice subjected to MCAO showed an increase in both parameters compared to sham mice. In the peri-infarct region, Iba-1^+^ cell showed a ramified morphology with an increase in the number of branch points (6.9±0.1, p≤0.001) and cellular area (1.03x10^6^±2.8x10^4^ µm^2^, p≤0.001) compared to the infarct zone (Figure 5E-F). In the latter, Iba-1^+^ cells showed an ameboid morphology with a significant reduction in both parameters (Branch points: 4.2±0.25, p≤0.001. Cell area: 5.8x10^5^±2.6x10^4^ µm^2^, p≤0.001) (Figure 5E-F).

Rapamycin-treated ischemic mice showed a significant decrease in both parameters compared to untreated ischemic counterparts. No differences were observed between the two damaged regions or between the times of rapamycin administration (Figure 5E-F). Slightly branched morphologies were observed compared to highly ramified morphologies MCAO in the peri-infarct region.

Astrocytes are another cell type involved in the inflammatory response. Analysis of the astrocytic response revealed that MCAO significantly increased the levels of GFAP (165.1±11.4%, p≤0.001) compared to the sham condition. The levels of GFAP in rapamycin-treated ischemic mice (pre- or post-MCAO) did not differ from those of untreated ischemic (Supplementary Figure 2A-C).

### mTORC1 inhibition ameliorates microglia reactivity after MCAO

To determine microglia reactivity after MCAO, we analysed soluble-TREM2 (sTREM2) and iNOS. The results refer to the entire region of damaged cerebral cortex because we did not find differences between the infarct zone and the peri-infarct region (data not shown). sTREM2 has been related to microglia activation in various neuroinflammation backgrounds (Moore et al., 2021). WB analysis showed an increase in sTREM2 levels after MCAO compared to untreated sham mice (140.2±8.4%, p≤0.001) (Figure 6A-B). Rapamycin-treated ischemic mice showed a significant reduction in sTREM2 levels compared to untreated ischemic (MCAO+R_pre_: 112.7±4.6%, p≤0.001; MCAO+R_post_: 87.20±8.3%, p≤0.05) (Figure 6A-B). This decrease was more marked when rapamycin was injected after MCAO induction. Rapamycin administration had no effect on the sham group (Figure 6A-B). In addition, we performed double immunofluorescence staining using Iba-1 and sTREM2 antibodies. We observed that MCAO increased the number of sTREM2^+^/Iba-1^+^ cells compared to the control (Figure 6C). The two ischemic groups treated with rapamycin showed a reduction in the number of sTREM2^+^/Iba-1^+^ cells compared to untreated counterparts (Figure 6C). In sham mice, rapamycin administration did not alter the number of sTREM2^+^/Iba-1^+^ cells (Figure 6C).

**Figure 6.**
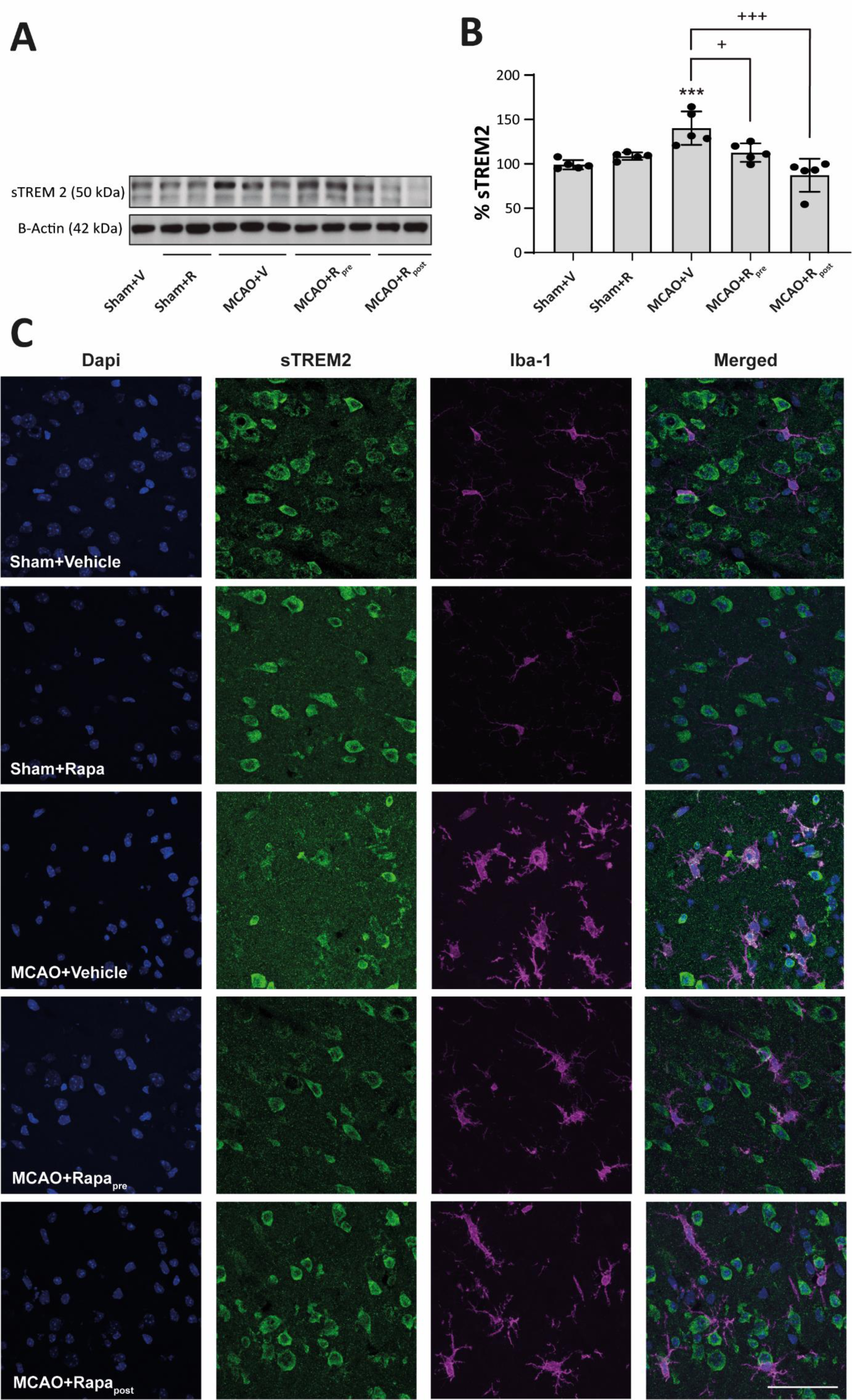
mTORC1 inhibition reduces Iba-1^+^/sTREM2^+^ after MCAO. A) Representative Western Blot images of the analysed extracts proteins from mice cerebral cortex. Immunodetections were performed using antibodies against sTREM2 and B-Actin, as a loading control. B) Graphical quantification of immunodetection by Western Blot. Data are normalized against B-Actin and expressed as the percentage of variation versus sham. Graph values represent means ± SEM (Ordinary one-way ANOVA *p ≤ 0.05; **p ≤ 0.01; ***p ≤ 0.001. Unparied t-test ^+^p ≤ 0.05; ^++^p ≤ 0.01; ^+++^p ≤ 0.001). Sham+V n=5, Sham+R n=5, MCAO+V n=5; MCAO+Rpre n=5 and MCAO+R_post_ n=5 (V=vehicle; R=rapamycin). * vs Sham+V and ^+^ vs MCAO+V group. C) Representative confocal immunofluorescence images from the entire damage area. Staining was performed using Dapi (blue), sTREM2 (green) and Iba-1 (magenta) antibodies. Scale bar: 25 µm.

To identify the effect of rapamycin on the abundance of microglia with the pro-inflammatory phenotype, we performed double immunofluorescence staining using antibodies against iNOS and Iba-1. Rapamycin administration did not alter the number of double-positive cells in sham mice. However, ischemic mice showed an increase in the number of iNOS^+^/Iba-1^+^ cells compared to untreated sham mice (Figure 7A). Rapamycin administration both pre and post-MCAO restored this parameter to sham values (Figure 7A). No differences were observed between rapamycin administration pre and post-MCAO (Figure 7A).

**Figure 7.**
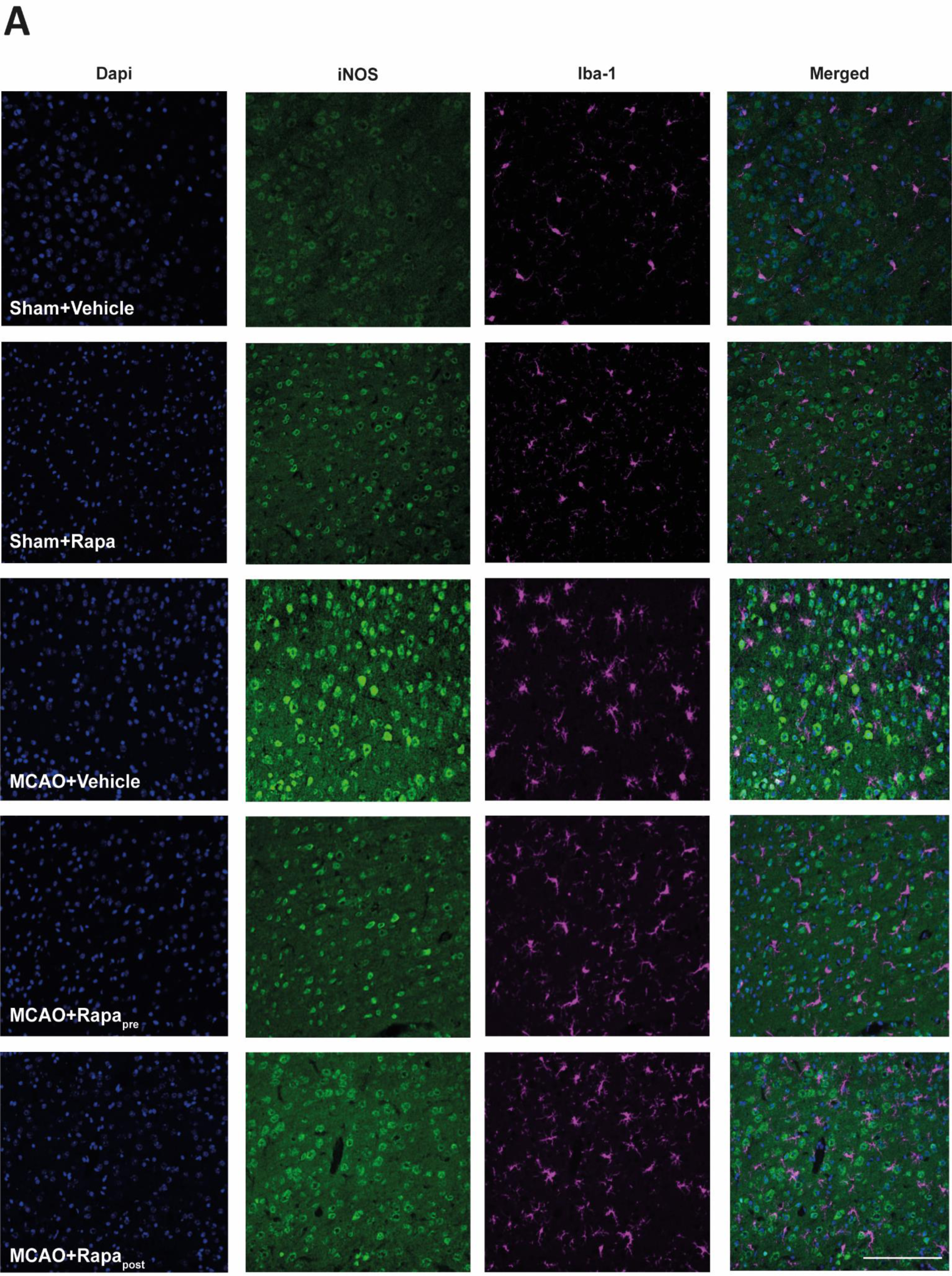
mTORC1 inhibition reduces Iba-1^+^/iNOS^+^ after MCAO. A) Representative confocal immunofluorescence images from the entire damage area. Staining was performed using Dapi (blue), iNOS (green) and Iba-1 (magenta) antibodies. Scale bar: 50 µm.

These results strongly support the notion that mTORC1 inhibition by rapamycin, both pre and post-MCAO, reduces pro-inflammatory microglia reactivity.

## Discussion

Ischemic stroke is a complex and multifactorial neurodegenerative disease associated with a large inflammatory process (Perez-Alvarez & Wandosell, 2016; Virani et al., 2021). Several authors support the notion that neuroinflammation plays a key role in the recovery of CNS homeostasis by modulating some brain repair mechanisms (Meschia & Brott, 2018). However, a sustained response over time may contribute to the development of psychiatric disorders and neurodegenerative diseases (Simats & Liesz, 2022; Xia et al., 2022). The therapeutic approaches used to reduce ischemic damage involve recanalization strategies, including the administration of antithrombotic agents and surgical removal of the thrombus to rescue less damage neurons in the peri-infarct area. The restoration of blood flow can mitigate the effects of ischemia only when performed a few hours after onset of the first symptoms. However, despite the beneficial effects of this approach, it is not suitable for all patients and does not reduce subsequent neurodegeneration (Sommer, 2017; Villa-González et al., 2022). In this context, considerable research efforts focus on the search for pharmacological alternatives to improve ischemic damage in these patients and enhance the effects of reperfusion.

Here we used a murine model of transient focal cerebral ischemia by intra-vasal thrombin injection, which has proven to be a suitable model to analyse the acute phase of ischemia in which neuroinflammation has been reported (Orset et al., 2007). As a therapeutic approach, we tested rapamycin, a known inhibitor of mTORC1, administered pre- and post-MCAO.

A pivotal player in the PI3K/Akt/mTORC1 survival pathway, the mTOR protein is associated with neuroprotection in the context of cerebral ischemia through its role in the maintenance of neuronal homeostasis (P. L. Fan et al., 2023). However, there is less information about its possible role in glial cells such as astrocytes and microglia. Recent studies suggest that the PI3K/Akt/mTORC1 pathway is involved in microglia activation and that the dysregulation of this pathway contributes to alterations of the neuroinflammatory response (Chu et al., 2021).

Our MRI data indicated that MCAO resulted in a considerable volume of lesion (18 mm^3^) in the parietal cortex, accompanied by a marked decrease in the number of neurons. The latter decrease was more pronounced in the infarct zone than in the peri-infarct region, as reported by others (Ermine et al., 2021). Cerebral ischemia triggers the early activation of glial cells, including microglia and astrocytes. Although the astrocyte response is later than the microglial response, in our model we detected an increase in GFAP levels distributed around the damaged area, thereby providing evidence of a response by these cells. Consistent with previous reports describing the microglial response as one of the earliest events after cerebral ischemia, we observed an increase in both the number of Iba-1^+^ cells and their stained area, thereby confirming microglial reactivity throughout the entire damaged area (Rupalla et al., 1998). In parallel, the biochemical analysis revealed an increase in the phosphorylation of mTOR and P70S6K, without any variations in their total amounts. Moreover, this analysis did not detect changes in either the phosphorylation or total levels of Akt. These data allow us to confirm that MCAO drives an increase in mTORC1 activity without affecting the basal activity of mTORC2, as reported elsewhere (Liang et al., 2018; Yang et al., 2015). However, other studies have described opposite results with respect to mTORC1 activity after ischemia (Chi et al., 2021; Gu et al., 2021; Mateos et al., 2016). The use of distinct models that included or excluded reperfusion, the duration of ischemic damage, and the time of analysis could explain the differences between the findings. In this context, to better understand the tissular response to ischemic damage and optimize strategies based on mTORC as therapeutic targets, it is important to take into account the model of ischemia used (Sommer, 2017).

The activation of microglia has been associated with changes in the morphology and functions of this cell type. These transformations require a balance between catabolic and anabolic cellular processes, which are regulated, in part, by mTORC1 (Chu et al., 2021). Thus, using an immunofluorescence approach, we revealed an increase in the number of p-S6^+^/Iba-1^+^ cells in the whole damaged area of MCAO mice. This observation strongly supports the notion that the increase in mTORC1 activity, observed by WB, is due mainly to microglia. In addition, the morphological heterogeneity of the microglial response has been associated with the extent and duration of ischemic damage, and with the characteristics of the brain region monitored by these cells (Masuda et al., 2011; Morrison & Filosa, 2013). Our data showed that the predominant microglial phenotype after MCAO differed between the infarct zone and peri-infarct region, with amoeboid microglia being most abundant in the former and highly branched microglia in the latter, as reported previously (Qin et al., 2019). Furthermore, microglia activation involves polarization to the M1 pro-inflammatory or M2 anti-inflammatory phenotype. Despite this binary classification, there is evidence that supports the coexistence of a heterogeneous activated microglia population (Girard et al., 2013; Kanazawa et al., 2017). In the present study, we analysed microglia reactivity using sTREM2 and iNOS. The former is a transmembrane receptor expressed mainly by immune cells and active microglia in the CNS (Moore et al., 2021). Some authors have reported an association between sTREM2 levels and the prognosis of ischemia (Kwon et al., 2020; Lu et al., 2022). On the other hand, iNOS produces nitric oxide, which promotes neuronal death after stroke (Tewari et al., 2021). Our data showed that MCAO triggers an increase in both markers in microglia, accompanied by an increase in the number of p-S6^+^/Iba-1^+^ in the damaged area. Based on our findings, we propose that, post-MCAO, mTORC1 activity mediates the early microglial activation that contributes to the detrimental effects of inflammation.

Thus, we studied whether mTORC1 inhibition confers neuroprotection in our ischemia model. Rapamycin has been used in clinical practice for several decades due to its immunosuppressive properties, and it shows promising results in some ischemic pathologies (Hadley et al., 2019; McKeage et al., 2003). However, there are controversial results about dosage and timing of administration. Several studies derived from meta-analysis have shown that low doses of rapamycin have greater efficacy against ischemic damage compared to high ones, showing a reduction in the volume of damage and neurological improvement in animals (Beard et al., 2019). Whereas mTORC1 is highly sensitive to rapamycin, mTORC2 inhibition requires higher doses or prolonged exposure to this drug (Fletcher et al., 2013; Sarbassov et al., 2005). Therefore, high rapamycin dosages exacerbate autophagy induction and mTORC2 inhibition. The regulation of neuronal survival pathways is under mTORC2 governance. Therefore, the maintenance of basal levels of this complex and/or its increase after cerebral ischemia could be a fundamental approach for neuronal survival and the reduction of the volume of damage (Beard et al., 2019; Kim et al., 2017). In this context, the inhibition of both mTORC1 and mTORC2 by rapalink-1 increases the volume of the ischemic lesion (Chi et al., 2016; Chi et al., 2021). In contrast, our results showed that 20 mg/kg rapamycin administered pre- or post-MCAO decreased the volume of damage by 35–45%. Our WB data showed that rapamycin did not modify Akt p-Ser473 levels in the sham group, thereby suggesting that this dosage of rapamycin is not sufficient to inhibit mTORC2 activity. In contrast, we observed that rapamycin caused a significant reduction in p-mTOR and p-P70S6K levels after MCAO—findings consistent with those reported in other studies (Liu et al., 2016; Yang et al., 2015).

Regarding the time of rapamycin administration, the present study and another carried out by Wu and colleagues (Wu et al., 2018) are the only ones to compare rapamycin administration pre- and post-ischemic damage. Our data showed that, for both rapamycin administration approaches, damage volume and neuronal loss were markedly lower compared to the vehicle-treated ischemic group, with no difference between the administration times used (Wu et al., 2018). However, at the biochemical level, rapamycin administration post-damage further reduced mTORC1 activity than at 48 h before ischemia, as reflected by lower levels of P70S6K phosphorylation by mTORC1 in the MCAO+R_post_ group. The differences observed could be attributed to the effective concentration of rapamycin in the brain, a parameter related to the time between drug administration and euthanasia of the animals.

mTORC1 inhibition led to a decrease in both the number of Iba-1^+^ cells and the intensity of Iba-1 staining in the damaged area, a finding consistent with other studies (Li et al., 2013). The analysis of microglia morphology showed that mTORC1 inhibition reduced the branching and area of Iba-1^+^ cells. Accordingly, after rapamycin treatment, we observed slightly branched morphologies in the damaged area compared to highly branched microglia in the peri-infarct region and amoeboid morphologies in the infarct zone described in untreated ischemic mice. The present study is the first to analyse the effects of rapamycin-induced mTORC1 inhibition on changes in microglial morphology. In addition, these morphological changes were associated with a significant reduction in sTREM2 and iNOS levels in microglia.

## Conclusions

On the basis of our results, we conclude that mTORC1 inhibition reduces the reactivity and the pro-inflammatory phenotype of microglia, thereby decreasing neuronal death and ischemic lesion volume. Several working hypotheses can be put forward to explain these findings. For instance, it is tempting to propose that the mTORC1-P70S6K-S6 pathway regulates microglia activation after the induction of cerebral ischemia and that the modulation of this activation by rapamycin reduces microglial reactivity and neuronal death. However, we cannot discard a complementary effect of rapamycin on neurons and /or other cells in the damaged area.

Finally, taken together, our data demonstrate that the use of specific mTORC1 inhibitors during the acute phase of cerebral ischemia has a neuroprotective effect, reducing the inflammatory microglial response. Given that rapamycin administration pre- and post-MCAO induction had similar effects with respect to neuronal protection, the administration of rapamycin or new rapalogs emerges as a potential therapeutic approach to tackle the acute phase of cerebral ischemia. Such a therapeutic strategy could be easily tested on patients as rapamycin is already authorised for the treatment of other human pathologies.

## List of abbreviations

Akt: Protein kinase B
CNS: Central Nervous System
iNOS: Inducible Nitric Oxide Synthase
MCAO: Middle Cerebral Artery Occlusion
MRI: Magnetic Resonance Imaging
mTOR: Mammalian target of rapamycin
mTORC1: Mammalian target of rapamycin complex 1
mTORC2: Mammalian target of rapamycin complex 2
P70S6K: Ribosomal protein S6 kinase
R: Rapamycin
S6: Ribosomal protein S6
sTREM2: Soluble Triggering receptor expressed on myeloid cells 2 (TREM)
V: Vehicle
WB: Western Blot

## Declarations

### Ethics approval and consent to participate

All the animals were housed in the Animal Facility of U1237 (GIP Cyceron) with food and water *ad libitum* access under a 12-hour light/dark cycle in a temperature-controlled environment. All care and experiments were performed following the ARRIVE guidelines (#24316) (www.nc3rs.org.uk).

### Consent for publication

Not applicable

### Availability of data and materials

The datasets used and/or analyses during the current study are available from the corresponding author on reasonable request.

### Authors’ contributions

**MVG:** data acquisition, data analysis, interpretation of data and drafted the manuscript. **MR:** data acquisition and discussion of results and revised manuscript. **GML:** data acquisition, data analysis. **PRM:** data acquisition and data analysis. **LVS:** data acquisition, data analysis. **DV:** conception and designing of the work and revised manuscript. **FW:** conception and designing of the work, interpretation of data, discussion of results and substantively revised manuscript. **MJP:** conception and designing of the work, interpretation of data, discussion of results and substantively revised manuscript.

## Acknowledgments

We would like to thank the Confocal Microscopy facility of the CBMSO for their assistance and support along this project. We are also thankful to the current and former members of our labs.

## References

Beard, D. J., Hadley, G., Thurley, N., Howells, D. W., Sutherland, B. A., & Buchan, A. M. (2019). The effect of rapamycin treatment on cerebral ischemia: A systematic review and meta-analysis of animal model studies. Int J Stroke, 14(2), 137–145. 10.1177/1747493018816503

Chauhan, A., Sharma, U., Jagannathan, N. R., & Gupta, Y. K. (2015). Rapamycin ameliorates brain metabolites alterations after transient focal ischemia in rats. Eur J Pharmacol, 757, 28–33. 10.1016/j.ejphar.2015.03.006

Chi, O. Z., Barsoum, S., Vega-Cotto, N. M., Jacinto, E., Liu, X., Mellender, S. J., & Weiss, H. R. (2016). Effects of rapamycin on cerebral oxygen supply and consumption during reperfusion after cerebral ischemia. Neuroscience, 316, 321–327. 10.1016/j.neuroscience.2015.12.045

Chi, O. Z., Liu, X., Cofano, S., Patel, N., Jacinto, E., & Weiss, H. R. (2021). Rapalink-1 Increased Infarct Size in Early Cerebral Ischemia-Reperfusion With Increased Blood-Brain Barrier Disruption. Front Physiol, 12, 706528. 10.3389/fphys.2021.706528

Chu, E., Mychasiuk, R., Hibbs, M. L., & Semple, B. D. (2021). Dysregulated phosphoinositide 3-kinase signaling in microglia: shaping chronic neuroinflammation. J Neuroinflammation, 18(1), 276. 10.1186/s12974-021-02325-6

Dirnagl, U., Iadecola, C., & Moskowitz, M. A. (1999). Pathobiology of ischaemic stroke: an integrated view. Trends Neurosci, 22(9), 391–397. 10.1016/s0166-2236(99)01401-0

Ermine, C. M., Bivard, A., Parsons, M. W., & Baron, J. C. (2021). The ischemic penumbra: From concept to reality. Int J Stroke, 16(5), 497–509. 10.1177/1747493020975229

Fan, J., Li, X., Yu, X., Liu, Z., Jiang, Y., Fang, Y., Zong, M., Suo, C., Man, Q., & Xiong, L. (2023). Global Burden, Risk Factor Analysis, and Prediction Study of Ischemic Stroke, 1990-2030. Neurology, 101(2), e137–e150. 10.1212/wnl.0000000000207387

Fan, P. L., Wang, S. S., Chu, S. F., & Chen, N. H. (2023). Time-dependent dual effect of microglia in ischemic stroke. Neurochem Int, 169, 105584. 10.1016/j.neuint.2023.105584

Fletcher, L., Evans, T. M., Watts, L. T., Jimenez, D. F., & Digicaylioglu, M. (2013). Rapamycin treatment improves neuron viability in an in vitro model of stroke. PLoS One, 8(7), e68281. 10.1371/journal.pone.0068281

Franklin, G. P. a. K. B. J. (2012). Paxinos and Franklin’s the Mouse Brain in Stereotaxic Coordinates.

Ge, P., Duan, H., Tao, C., Niu, S., Hu, Y., Duan, R., Shen, A., Sun, Y., & Sun, W. (2023). TMAO Promotes NLRP3 Inflammasome Activation of Microglia Aggravating Neurological Injury in Ischemic Stroke Through FTO/IGF2BP2. J Inflamm Res, 16, 3699–3714. 10.2147/jir.S399480

Girard, S., Brough, D., Lopez-Castejon, G., Giles, J., Rothwell, N. J., & Allan, S. M. (2013). Microglia and macrophages differentially modulate cell death after brain injury caused by oxygen-glucose deprivation in organotypic brain slices. Glia, 61(5), 813–824. 10.1002/glia.22478

Glass, C. K., Saijo, K., Winner, B., Marchetto, M. C., & Gage, F. H. (2010). Mechanisms underlying inflammation in neurodegeneration. Cell, 140(6), 918–934. 10.1016/j.cell.2010.02.016

Hadley, G., Beard, D. J., Couch, Y., Neuhaus, A. A., Adriaanse, B. A., DeLuca, G. C., Sutherland, B. A., & Buchan, A. M. (2019). Rapamycin in ischemic stroke: Old drug, new tricks? J Cereb Blood Flow Metab, 39(1), 20–35. 10.1177/0271678x18807309

Hernández, I. H., Villa-González, M., Martín, G., Soto, M., & Pérez-Álvarez, M. J. (2021). Glial Cells as Therapeutic Approaches in Brain Ischemia-Reperfusion Injury. Cells, 10(7). 10.3390/cells10071639

Jurcau, A., & Simion, A. (2021). Neuroinflammation in Cerebral Ischemia and Ischemia/Reperfusion Injuries: From Pathophysiology to Therapeutic Strategies. Int J Mol Sci, 23(1). 10.3390/ijms23010014

Kanazawa, M., Ninomiya, I., Hatakeyama, M., Takahashi, T., & Shimohata, T. (2017). Microglia and Monocytes/Macrophages Polarization Reveal Novel Therapeutic Mechanism against Stroke. Int J Mol Sci, 18(10). 10.3390/ijms18102135

Karalis, V., & Bateup, H. S. (2021). Current Approaches and Future Directions for the Treatment of mTORopathies. Dev Neurosci, 43(3-4), 143–158. 10.1159/000515672

Kim, Y. S., Yoo, A., Son, J. W., Kim, H. Y., Lee, Y. J., Hwang, S., Lee, K. Y., Lee, Y. J., Ayata, C., Kim, H. H., & Koh, S. H. (2017). Early Activation of Phosphatidylinositol 3-Kinase after Ischemic Stroke Reduces Infarct Volume and Improves Long-Term Behavior. Mol Neurobiol, 54(7), 5375–5384. 10.1007/s12035-016-0063-4

Kwon, H. S., Lee, E. H., Park, H. H., Jin, J. H., Choi, H., Lee, K. Y., Lee, Y. J., Lee, J. H., de Oliveira, F. M. S., Kim, H. Y., Seo Kim, Y., Kim, B. J., Heo, S. H., Chang, D. I., Kamali-Moghaddam, M., & Koh, S. H. (2020). Early increment of soluble triggering receptor expressed on myeloid cells 2 in plasma might be a predictor of poor outcome after ischemic stroke. J Clin Neurosci, 73, 215–218. 10.1016/j.jocn.2020.02.016

Leyh, J., Paeschke, S., Mages, B., Michalski, D., Nowicki, M., Bechmann, I., & Winter, K. (2021). Classification of Microglial Morphological Phenotypes Using Machine Learning. Front Cell Neurosci, 15, 701673. 10.3389/fncel.2021.701673

Li, D., Wang, C., Yao, Y., Chen, L., Liu, G., Zhang, R., Liu, Q., Shi, F. D., & Hao, J. (2016). mTORC1 pathway disruption ameliorates brain inflammation following stroke via a shift in microglia phenotype from M1 type to M2 type. Faseb j, 30(10), 3388–3399. 10.1096/fj.201600495R

Li, T., Pang, S., Yu, Y., Wu, X., Guo, J., & Zhang, S. (2013). Proliferation of parenchymal microglia is the main source of microgliosis after ischaemic stroke. Brain, 136(Pt 12), 3578–3588. 10.1093/brain/awt287

Liu, G. Y., & Sabatini, D. M. (2020). mTOR at the nexus of nutrition, growth, ageing and disease. Nat Rev Mol Cell Biol, 21(4), 183–203. 10.1038/s41580-019-0199-y

Liu, P., Yang, X., Hei, C., Meli, Y., Niu, J., Sun, T., & Li, P. A. (2016). Rapamycin Reduced Ischemic Brain Damage in Diabetic Animals Is Associated with Suppressions of mTOR and ERK1/2 Signaling. Int J Biol Sci, 12(8), 1032–1040. 10.7150/ijbs.15624

Liu, Y., Xue, X., Zhang, H., Che, X., Luo, J., Wang, P., Xu, J., Xing, Z., Yuan, L., Liu, Y., Fu, X., Su, D., Sun, S., Zhang, H., Wu, C., & Yang, J. (2019). Neuronal-targeted TFEB rescues dysfunction of the autophagy-lysosomal pathway and alleviates ischemic injury in permanent cerebral ischemia. Autophagy, 15(3), 493–509. 10.1080/15548627.2018.1531196

Lu, Y., Zhao, Y., Zhang, Q., Fang, C., Bao, A., Dong, W., Peng, Y., Peng, H., Ju, Z., He, J., Zhang, Y., Xu, T., & Zhong, C. (2022). Soluble TREM2 is associated with death and cardiovascular events after acute ischemic stroke: an observational study from CATIS. J Neuroinflammation, 19(1), 88. 10.1186/s12974-022-02440-y

Mansfield, A., Inness, E. L., & McIlroy, W. E. (2018). Stroke. Handb Clin Neurol, 159, 205–228. 10.1016/b978-0-444-63916-5.00013-6

Masuda, T., Croom, D., Hida, H., & Kirov, S. A. (2011). Capillary blood flow around microglial somata determines dynamics of microglial processes in ischemic conditions. Glia, 59(11), 1744–1753. 10.1002/glia.21220

Mateos, L., Perez-Alvarez, M. J., & Wandosell, F. (2016). Angiotensin II type-2 receptor stimulation induces neuronal VEGF synthesis after cerebral ischemia. Biochim Biophys Acta, 1862(7), 1297–1308. 10.1016/j.bbadis.2016.03.013

McKeage, K., Murdoch, D., & Goa, K. L. (2003). The sirolimus-eluting stent: a review of its use in the treatment of coronary artery disease. Am J Cardiovasc Drugs, 3(3), 211–230. 10.2165/00129784-200303030-00007

Meschia, J. F., & Brott, T. (2018). Ischaemic stroke. Eur J Neurol, 25(1), 35–40. 10.1111/ene.13409

Meyuhas, O. (2015). Ribosomal Protein S6 Phosphorylation: Four Decades of Research. Int Rev Cell Mol Biol, 320, 41–73. 10.1016/bs.ircmb.2015.07.006

Moore, E. E., Liu, D., Li, J., Schimmel, S. J., Cambronero, F. E., Terry, J. G., Nair, S., Pechman, K. R., Moore, M. E., Bell, S. P., Beckman, J. A., Gifford, K. A., Hohman, T. J., Blennow, K., Zetterberg, H., Carr, J. J., & Jefferson, A. L. (2021). Association of Aortic Stiffness With Biomarkers of Neuroinflammation, Synaptic Dysfunction, and Neurodegeneration. Neurology, 97(4), e329–e340. 10.1212/wnl.0000000000012257

Morrison, H. W., & Filosa, J. A. (2013). A quantitative spatiotemporal analysis of microglia morphology during ischemic stroke and reperfusion. J Neuroinflammation, 10, 4. 10.1186/1742-2094-10-4

Orset, C., Macrez, R., Young, A. R., Panthou, D., Angles-Cano, E., Maubert, E., Agin, V., & Vivien, D. (2007). Mouse model of in situ thromboembolic stroke and reperfusion. Stroke, 38(10), 2771–2778. 10.1161/strokeaha.107.487520

Pan, Y. W., Wu, D. P., Liang, H. F., Tang, G. Y., Fan, C. L., Shi, L., Ye, W. C., & Li, M. M. (2022). Total Saponins of Panax notoginseng Activate Akt/mTOR Pathway and Exhibit Neuroprotection in vitro and in vivo against Ischemic Damage. Chin J Integr Med, 28(5), 410–418. 10.1007/s11655-021-3454-y

Paolicelli, R. C., Sierra, A., Stevens, B., Tremblay, M. E., Aguzzi, A., Ajami, B., Amit, I., Audinat, E., Bechmann, I., Bennett, M., Bennett, F., Bessis, A., Biber, K., Bilbo, S., Blurton-Jones, M., Boddeke, E., Brites, D., Brône, B., Brown, G. C., … Wyss-Coray, T. (2022). Microglia states and nomenclature: A field at its crossroads. Neuron, 110(21), 3458–3483. 10.1016/j.neuron.2022.10.020

Perez-Alvarez, M. J., Mateos, L., Alonso, A., & Wandosell, F. (2015). Estradiol and Progesterone Administration After pMCAO Stimulates the Neurological Recovery and Reduces the Detrimental Effect of Ischemia Mainly in Hippocampus. Mol Neurobiol, 52(3), 1690–1703. 10.1007/s12035-014-8963-7

Pérez-Álvarez, M. J., Maza Mdel, C., Anton, M., Ordoñez, L., & Wandosell, F. (2012). Post-ischemic estradiol treatment reduced glial response and triggers distinct cortical and hippocampal signaling in a rat model of cerebral ischemia. J Neuroinflammation, 9, 157. 10.1186/1742-2094-9-157

Perez-Alvarez, M. J., Villa Gonzalez, M., Benito-Cuesta, I., & Wandosell, F. G. (2018). Role of mTORC1 Controlling Proteostasis after Brain Ischemia. Front Neurosci, 12, 60. 10.3389/fnins.2018.00060

Perez-Alvarez, M. J., & Wandosell, F. (2016). Stroke and Neuroinflamation: Role of Sexual Hormones. Curr Pharm Des, 22(10), 1334–1349. 10.2174/138161282210160304112834

Qin, C., Zhou, L. Q., Ma, X. T., Hu, Z. W., Yang, S., Chen, M., Bosco, D. B., Wu, L. J., & Tian, D. S. (2019). Dual Functions of Microglia in Ischemic Stroke. Neurosci Bull, 35(5), 921–933. 10.1007/s12264-019-00388-3

Rupalla, K., Allegrini, P. R., Sauer, D., & Wiessner, C. (1998). Time course of microglia activation and apoptosis in various brain regions after permanent focal cerebral ischemia in mice. Acta Neuropathol, 96(2), 172–178. 10.1007/s004010050878

Sarbassov, D. D., Guertin, D. A., Ali, S. M., & Sabatini, D. M. (2005). Phosphorylation and regulation of Akt/PKB by the rictor-mTOR complex. Science, 307(5712), 1098–1101. 10.1126/science.1106148

Simats, A., & Liesz, A. (2022). Systemic inflammation after stroke: implications for post-stroke comorbidities. EMBO Mol Med, 14(9), e16269. 10.15252/emmm.202216269

Sommer, C. J. (2017). Ischemic stroke: experimental models and reality. Acta Neuropathol, 133(2), 245–261. 10.1007/s00401-017-1667-0

Tewari, D., Sah, A. N., Bawari, S., Nabavi, S. F., Dehpour, A. R., Shirooie, S., Braidy, N., Fiebich, B. L., Vacca, R. A., & Nabavi, S. M. (2021). Role of Nitric Oxide in Neurodegeneration: Function, Regulation, and Inhibition. Curr Neuropharmacol, 19(2), 114–126. 10.2174/1570159x18666200429001549

Villa-González, M., Martín-López, G., & Pérez-Álvarez, M. J. (2022). Dysregulation of mTOR Signaling after Brain Ischemia. Int J Mol Sci, 23(5). 10.3390/ijms23052814

Virani, S. S., Alonso, A., Aparicio, H. J., Benjamin, E. J., Bittencourt, M. S., Callaway, C. W., Carson, A. P., Chamberlain, A. M., Cheng, S., Delling, F. N., Elkind, M. S. V., Evenson, K. R., Ferguson, J. F., Gupta, D. K., Khan, S. S., Kissela, B. M., Knutson, K. L., Lee, C. D., Lewis, T. T., … Tsao, C. W. (2021). Heart Disease and Stroke Statistics-2021 Update: A Report From the American Heart Association. Circulation, 143(8), e254–e743. 10.1161/cir.0000000000000950

Wu, M., Zhang, H., Kai, J., Zhu, F., Dong, J., Xu, Z., Wong, M., & Zeng, L. H. (2018). Rapamycin prevents cerebral stroke by modulating apoptosis and autophagy in penumbra in rats. Ann Clin Transl Neurol, 5(2), 138–146. 10.1002/acn3.507

Xia, Q., Zhan, G., Mao, M., Zhao, Y., & Li, X. (2022). TRIM45 causes neuronal damage by aggravating microglia-mediated neuroinflammation upon cerebral ischemia and reperfusion injury. Exp Mol Med, 54(2), 180–193. 10.1038/s12276-022-00734-y

Yang, X., Hei, C., Liu, P., Song, Y., Thomas, T., Tshimanga, S., Wang, F., Niu, J., Sun, T., & Li, P. A. (2015). Inhibition of mTOR Pathway by Rapamycin Reduces Brain Damage in Rats Subjected to Transient Forebrain Ischemia. Int J Biol Sci, 11(12), 1424–1435. 10.7150/ijbs.12930

Zietz, A., Gorey, S., Kelly, P. J., Katan, M., & McCabe, J. (2023). Targeting inflammation to reduce recurrent stroke. Int J Stroke, 17474930231207777. 10.1177/17474930231207777

