## Supplementary figures and images for "Pharmacological inhibition of mTORC1 reduces neural death and damage volume after MCAO by modulating microglial reactivity"

### Supplemental Figure 1

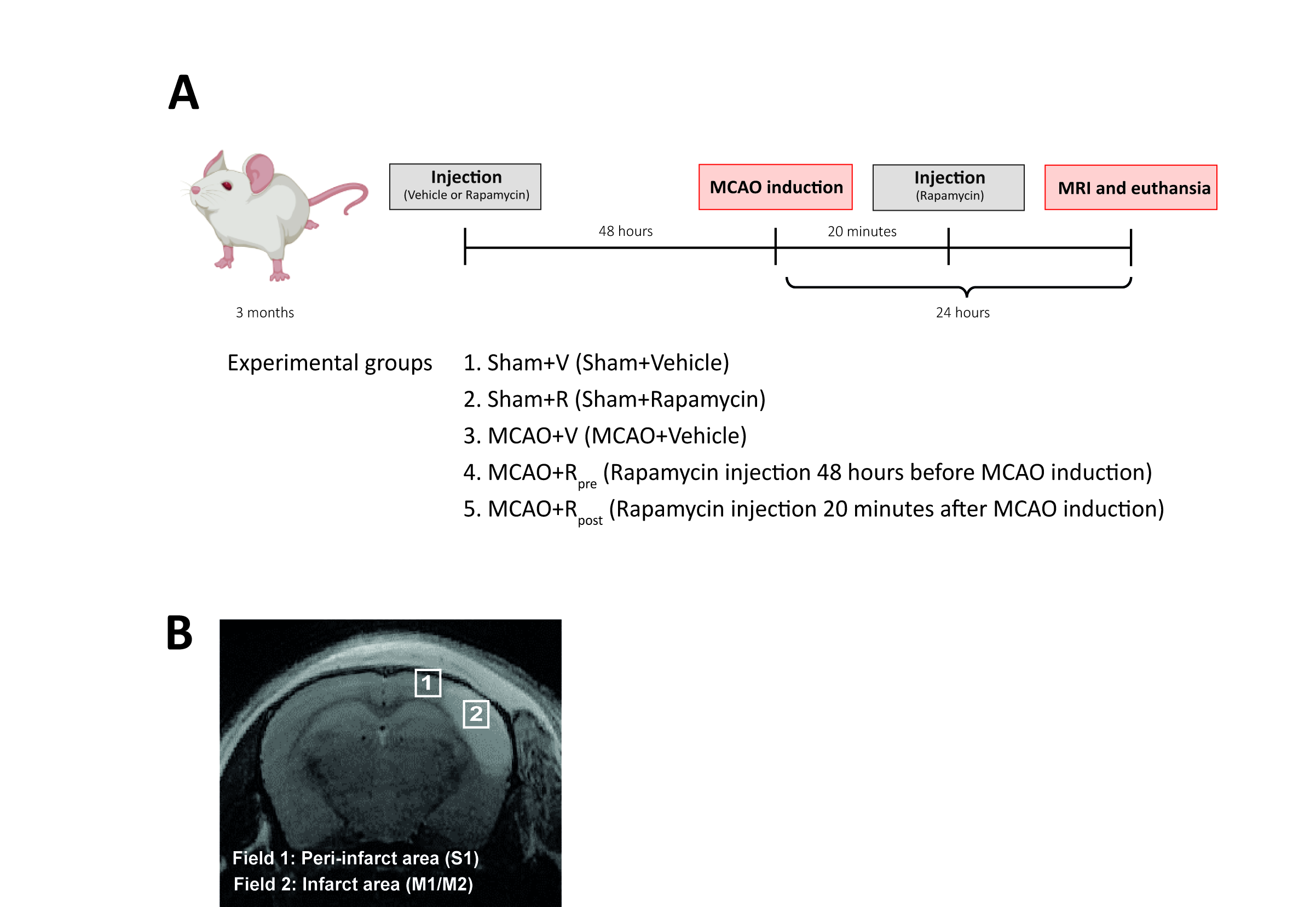

### Supplemental Figure 2

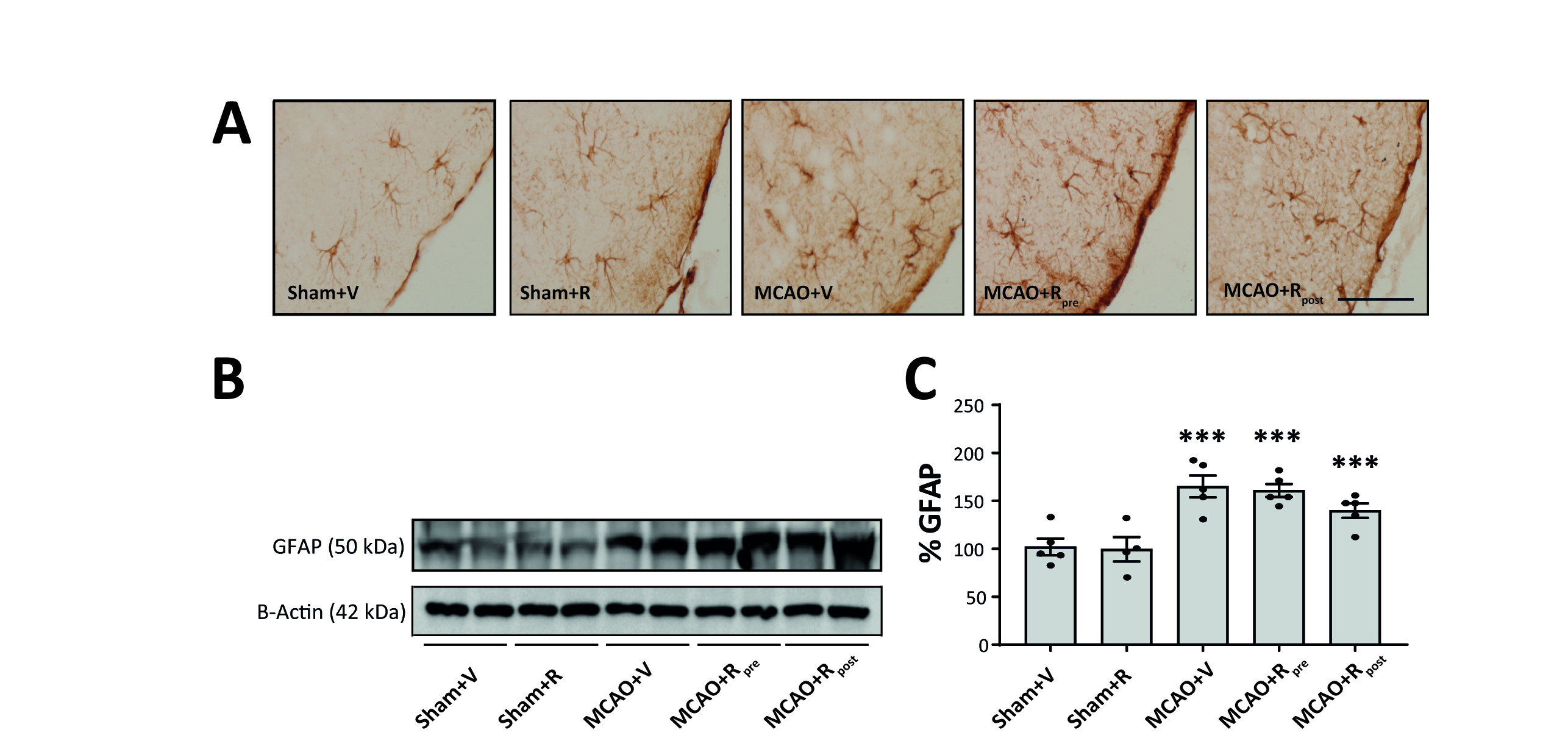
